# Prevalence of Congenital Amusia

**DOI:** 10.1101/070961

**Authors:** Isabelle Peretz, Dominique T. Vuvan

## Abstract

Congenital amusia (commonly known as tone-deafness) is a lifelong musical disorder that should affect 4% of the population according to a single estimate based on a single test from 1980. Here we present the first large-based measure of prevalence with a sample of 20,000 participants that does not rely on self-referral. On the basis of three objective tests and a questionnaire, we show that (a) the prevalence of congenital amusia is only 1.5% with slightly more females than males, unlike other developmental disorders where males often predominate; (b) self-disclosure is a reliable index of congenital amusia, that suggests that congenital amusia is hereditary with 46% first-degree relatives similarly affected; c) that the deficit is not attenuated by musical training and d) it emerges in relative isolation from other cognitive disorder except for spatial orientation problems. Hence, we suggest that congenital amusia is likely to result from genetic variations that affect musical abilities specifically.

## Introduction

Music engagement is a fundamental human trait. The study of its brain basis has been increasingly scrutinized and advances in molecular technologies make the genomic factors that are associated with its emergence possible. The search for the genetic correlates of musicality has gained interest ^1^ but it still lags behind research in other cognitive domains such as language. In language, the study of speech disorders has been instrumental in the identification of the underlying genes, namely FOXP2 (ref. 2). Likewise, the study of specific musical disorders, referred to as congenital amusia, an umbrella term for lifelong musical disabilities, may provide new entry points for deciphering the key neurobiological pathways for music. Indeed, congenital amusia results from a neuronal anomaly affecting the right auditory cortex and its connection to the inferior frontal gyrus.^3–7^ Congenital amusia also appears hereditary.^8^ Thus, congenital amusia represents a unique opportunity to trace the causal links between music, brain and genes.

Amusia is an accident of nature that affects musical abilities. Traditionally, amusia referred to a failure to process music as a consequence of brain damage.^9^ More recently, a congenital form of amusia with no history of brain injury (MIM 191200) has been uncovered.^10^ Individuals with the most common form of congenital amusia have a normal understanding of speech. They can recognize speakers by their voices and can identify all sorts of familiar environmental sounds such as animal cries. What distinguishes them from ordinary people is their difficulty with recognizing a familiar tune without the aid of the lyrics, and their inability to detect when someone sings out-of-tune, including themselves. Most notably, amusics fail to detect “wrong notes” (off-key notes) in conventional but unfamiliar melodies.^11^ This behavioral failure is diagnostic since there is no overlap between the distributions of the scores of amusics and controls. Thus, this musical pitch disorder presents a clear-cut phenotype that calls for genetic analyses.

Evidence for the notion that musical pitch processing might be a good target for phenotype-genotype correlations comes from a family aggregation study of amusia by our group^8^ and from an independent twin study.^12^ In the family aggregation study, the incidence of amusia was quantified by auditory testing of 71 members of 9 large families of amusic probands, as well as of 75 members of 10 control families. The musical pitch disorder was expressed in 39% of first-degree relatives in amusic families whereas it was only present in 3% in control families. This incidence of amusia is of the same order of magnitude as the heritability of speech disorders.^13^

In the twin study, monozygotic and dizygotic pairs were required to detect anomalous pitches in popular melodies (using the Distorted Tune Test; DTT). Genetic model-fitting indicated that the influence of shared genes was more important than shared environments, with a heritability of 70-80%. The DTT has been administered to more than 600 participants in the U.K.^14^ Approximately 4% of this sample performed as poorly as 20 adults who identified themselves or were identified by others as amusic. This suggests that 4% of the population may suffer from a genetically determined defect in perceiving musical pitch. However, both the twin and prevalence study relied on a single measurement of musical pitch ability (the DTT) that has poor sensitivity and validity. On this single test, the majority (78.5%) of participants achieved a perfect score. Moreover, the test uses well-known melodies and has no control condition. Therefore, it remains unclear if amusics who fail on the DTT do so because of a specific deficit in melody perception or because of a more general problem such as a memory or attention deficit. The measures used in the family aggregation study of congenital amusia^8^ have neither of these shortcomings and were used here to establish the prevalence of congenital amusia.

Congenital amusia was established here with the help of three tests. The first corresponds to the Scale test of the Montreal Battery of Evaluation of Amusia (MBEA^15^) because this is the most widely used screening test for amusia.^16^ The Scale test consists of comparing 30 pairs of melodies that differ by an off-key note in half of the trials. The other two tests, the Off-key and Off-beat tests, include the detection of either an off-key or an out-of-time note in the same melodies. The Off-beat test serves as a control condition because the most common (pitch-based) form of congenital amusia does not typically affect performance on this test.^8^ Additionally, the Off-beat test may serve as an indication for the presence of a new form of congenital amusia that is not pitch-based but time-based, and to which we refer as beat deafness.^17^ These three tests, the Scale, Off-key and Off-beat test, present a number of advantages over the DTT, the test that served to provide the first and only estimate of the prevalence of congenital amusia. First, the use of unfamiliar melodies makes these tests relatively culture-free. For example, these tests have been effective in identifying cases of congenital amusia among speakers of tone languages.^18^ Second, the use of the off-beat test as a control condition allowed us to exclude participants with non-musical problems such as attention deficits. Third, these tests have been validated with the full MBEA battery.^19^ Finally, these tests have been available on the internet for nearly 10 years and have reached a large and diverse population. In sum, these tests currently constitute the best available tool to revisit the prevalence of congenital amusia.

Reliable measures of prevalence are useful to narrow down the responsible genes for the disorder and the identification of factors that are associated to its expression. For instance, neurodevelopmental disorders such as autism (MIM 209850) affect more males than females. Congenital amusia may also be associated with other developmental disorders, such as dyslexia (MIM 127700), because both disorders seem to result from anomalous neuronal proliferation and migration in the auditory cortex (ectopias^4^). However, musical training may attenuate the expression of the disorder. All these parameters may interact to make estimating prevalence a complex task that can only be addressed by a large population survey.

## METHOD

Participants volunteered to test their musical abilities via our website (www.brams.umontreal.ca/amusia-general) between July 2008 and December 2015. They could choose their language of preference (English or French) and complete the three tests of musical ability, the Scale, Off-beat and Off-key tests, followed by a self-report inventory. If taken without pauses, the entire procedure takes about 30 minutes. Participants were then given online feedback on their scores on the three musical tests. There was no incentive for the participants other than their individual scores provided at the end of the survey.

### Participants

From the 20,850 who participated, we analyzed the data of 16,625 adults (aged 18 – 65; Mean age was 31.6, *SD*=12.8) without reported history of head trauma and hearing loss, who completed all 3 tests without repeat and who detected the catch trial inserted near the end of the first Scale test (Figure 1). The catch was a comparison melody in which an obvious pitch change was embedded.

About half (50.6%) the volunteers were females. They were mostly tested in English (67.0%). The sample contained a large spread in terms of education (still in education: 5%, secondary school degree: 18%, undergraduate degree/professional qualification: 45%, postgraduate degree: 32%). Nevertheless, the current sample was relatively educated with a mean number of years of education of 16.9 (SD = 3.9). Less than 2% (1.4%) stated ‘Music’ as their occupation.

**Figure 1.**
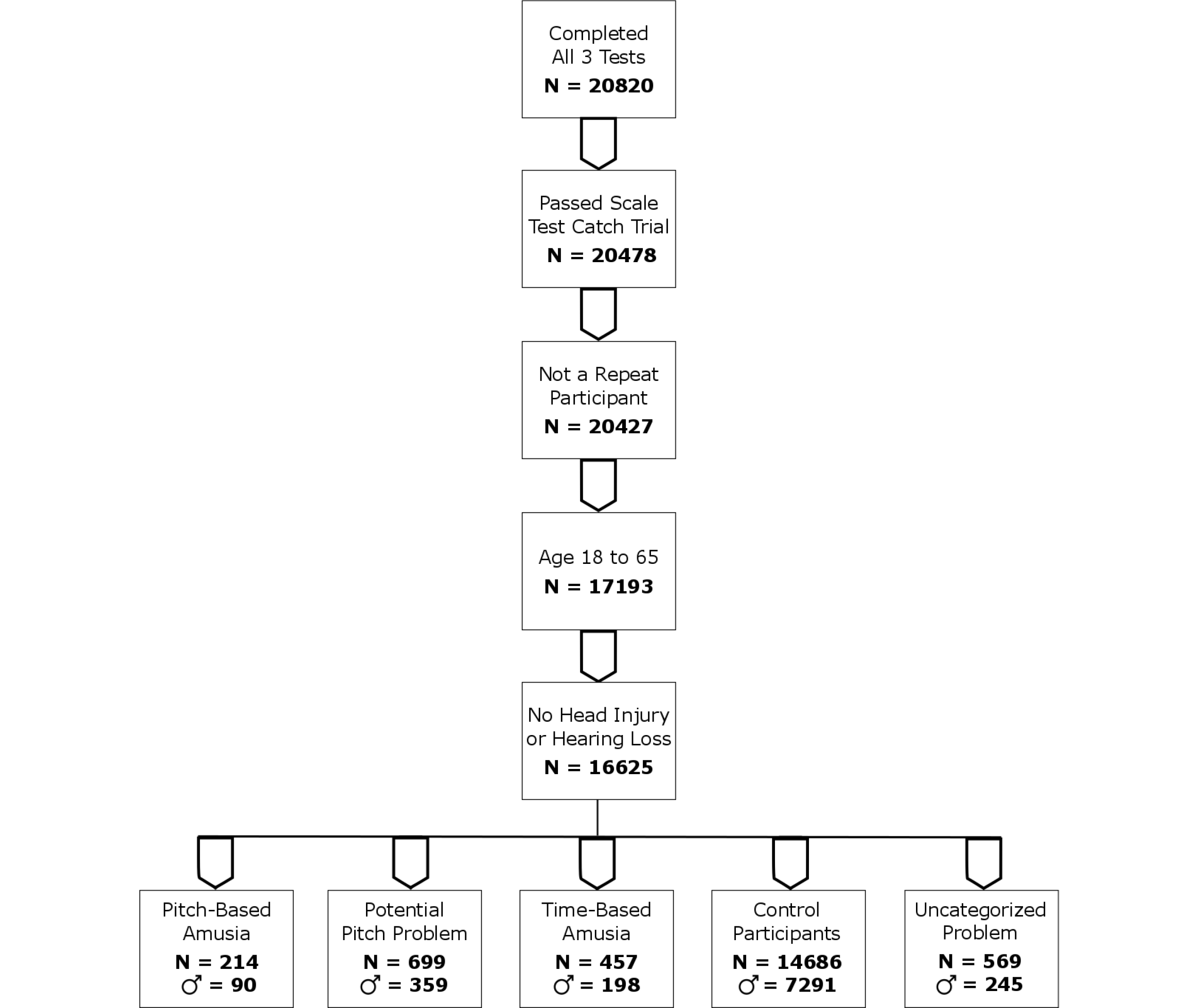
Data filtering procedure. Potential Pitch Problem means that participants failed only one of the two pitch tests. Uncategorized Problem refers to participants who failed both the off-beat test and at least one pitch test.

### Procedure

The entire assessment took place online. Participants gave informed consent before completing the survey. They were then prompted to test and adjust the volume of their audio equipment before starting the *scale* test of the Montreal Battery for the Evaluation of Amusia.^15^ As mentioned, it comprises 30 pairs of melodies, presented with a piano timbre. In half of the pairs, a key-violated alternate melody modifies the pitch of one tone so that it is out of key while maintaining the original melodic contour. In the other half, the melody is repeated without change. Participants’ task is to determine whether the two melodies are the same or not. A catch trial involved an alternate melody in which one full measure contains random pitches over several octaves. The second task is the *off-beat* condition and the third task is the *off-key* condition. In both conditions, half the melodies are congruous. In the other half, the same critical tone is altered either in time or in pitch. The critical tone always falls on the first downbeat in the third bar of the four-bar melody (hence, is metrically stressed) and is 500 ms long. The time change of the *off-beat* condition consists of introducing a silence of 5/7 of the beat duration (i.e., 357 ms) directly preceding the critical tone, thereby locally disrupting the meter (i.e., regularity). In the *off-key* condition, the change consists of using a tone that is outside the key of the melody, hence introducing a “foreign” or “wrong” note (Figure 2). The melodies are presented with 10 different timbres (e.g., piano, saxophone, clarinet, recorder, harp, strings, guitar) in order to make the auditory test more engaging. In each condition, subjects are presented with 24 melodies (12 congruous, 12 incongruous) one at a time, in a random order. Their task is simply to detect whether an incongruity occurs in each melody, by way of clicking a “yes” button whenever they detect an anomaly, and a “no” button when they do not detect an incongruity. Participants receive 2 practice trials before each test and are provided with feedback after each practice trial.

**Figure 2.**
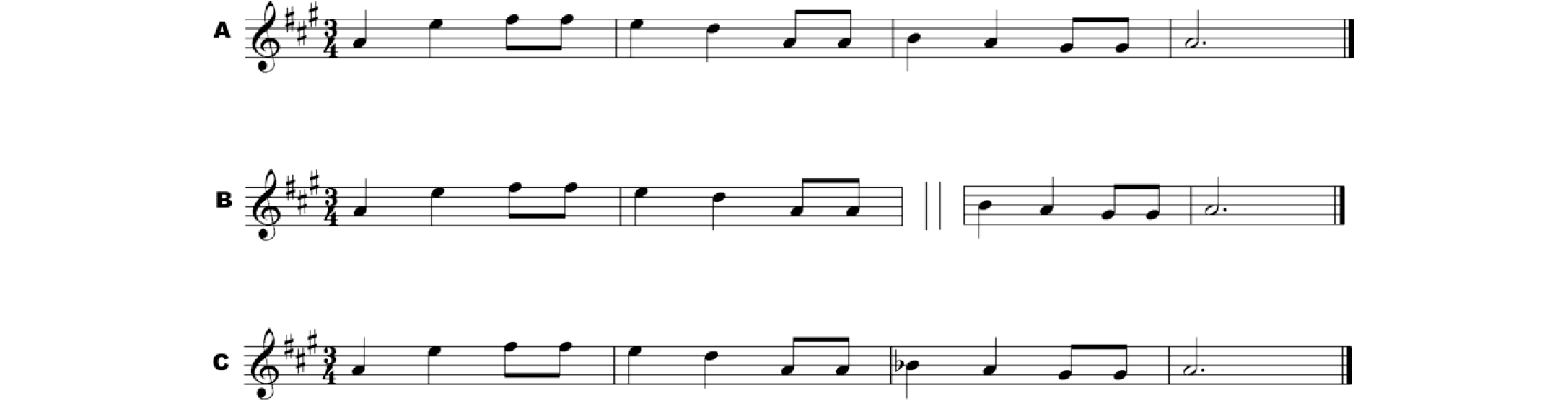
Example of stimuli: (A) a melody with no manipulation; (B) a melody with a temporal anomaly (Off-beat test) and (C) a melody with a pitch anomaly (Scale and Off-key tests).

The three tests use the same set of unfamiliar melodies constructed according to Western music tonal conventions and computer-generated at a tempo of 120 beats/min. The stimuli are presented in MP3 using the open source LAME MP3 encoding package and standard Web browser technologies (i.e., HTML, PHP, and Flash). The responses are fully anonymised, automatically recorded and tabulated for further analysis in Microsoft Excel.

## RESULTS AND COMMENTS

For each test, we used the standard criterion of 2 *SD* below the mean as a cut-off below which scores were indicative of a disorder (Table 1). Furthermore, to control for motivation or attentional problems, we selected participants who scored above the cut-off in the Off-beat test and below cut-off on both the Scale and Off-key tests for pitch-based amusia (n=214, 57.9% females). Conversely, for time-based amusia, we considered scores above the cut-off in the Scale and Off-key tests but below cut-off in the Off-beat test (n=457, 56.7% females). Controls scored above cut-off in all three tests (n=14,686, 50.3 % females). Because a pitch-based deficit could affect the Scale test or the Off-key test, scores below cut-off on both tests was considered as indicative of amusia (prevalence for each test can be found in Supplementary Table 1). Performance for each resulting group on each test is presented in Table 1. There was no performance difference between female and male participants in any of the tests and for none of the groups considered.

**Table 1.** Group performance on each of the three conditions assessed

| Tests | Controls<br>(N= 14,686 ) | Pitch-Based Amusics<br>(N= 214) | Time-Based Amusics<br>(N= 457) |
| --- | --- | --- | --- |
| Out-of- Scale (/30) | 26.65 (2.38) | 17.20 (1.78)* | 25.59 (2.70)* |
| Off-key (/24) | 20.85 (2.29) | 11.70 (1.27)* | 19.15 (2.54)* |
| Off-beat (/24) | 20.43 (1.87) | 18.92 (1.76)* | 13.86 (1.60)* |

*Note.* \* indicates that the score is significantly different from controls at $p < .05$ .

The scores obtained by controls in the two pitch (Scale and Off-key) tests were correlated as expected, with r(14899) = .45, p < .001. Less expected was the finding of a significant correlation between the Off-beat and Off-key tests, r(14899) = .25, p < .001.

### Prevalence of amusia

By the conservative criteria used for pitch-based amusia, 1.5% of the population qualifies for having the disorder. It is slightly higher among women (57.9%) than men (42.1%), χ^2^(1)= 4.87, p = .03. This difference (according to self-reported gender) could not be accounted by differences in age nor by regular or music education (all p > .05).

### Musical training

Nearly half (48%) of the pitch-based amusics reported no musical training besides mandatory music classes at school while only 25 % of controls reported no training. Moreover, the 52% of the amusics who did get extra-curricular lessons did so for a shorter duration but it is by no means negligible (Table 2). Having extra-curricular lessons does not seem to affect the severity of the disorder, nor performance in general.

Correlations between years of musical education and performance on each test are weak in general (all *r* values below .15).

**Table 2.**
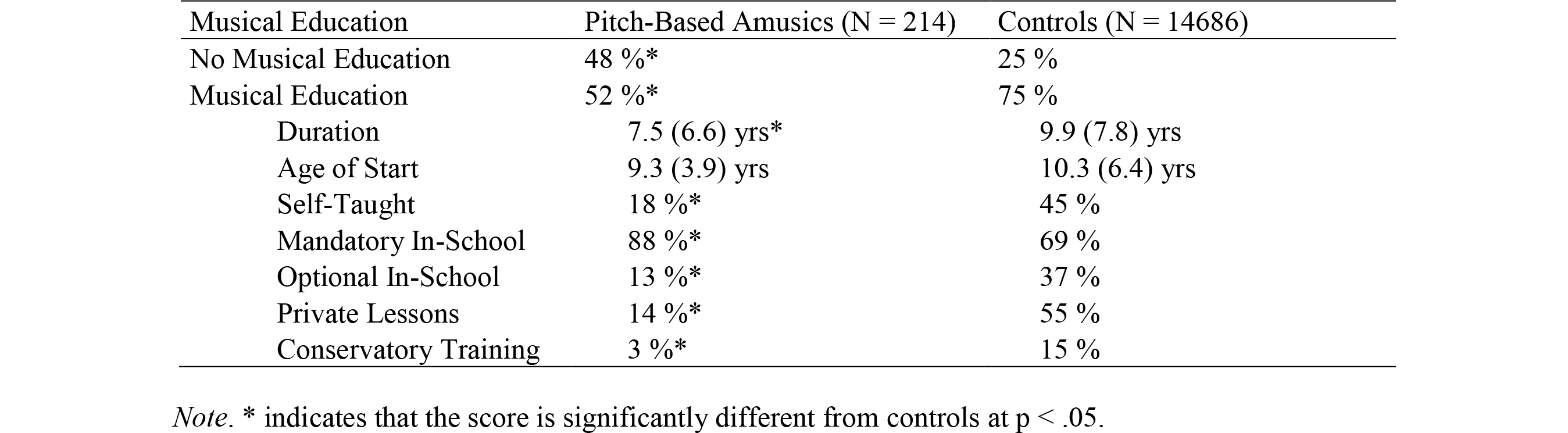
Musical education details for control participants and pitch-based amusics

| Musical Education | Pitch-Based Amusics (N = 214) | Controls (N = 14686) |
| --- | --- | --- |
| No Musical Education | 48 % * | 25 % |
| Musical Education | 52 % * | 75 % |
| Duration | 7.5 (6.6) yrs* | 9.9 (7.8) yrs |
| Age of Start | 9.3 (3.9) yrs | 10.3 (6.4) yrs |
| Self-Taught | 18 % * | 45 % |
| Mandatory In-School | 88 % * | 69 % |
| Optional In-School | 13 % * | 37 % |
| Private Lessons | 14 % * | 55 % |
| Conservatory Training | 3 % * | 15 % |

*Note.* \* indicates that the score is significantly different from controls at $p < .05$ .

**Self-report** of amusic problems is instrumental in research for there is no system to identify amusic cases in the regular curriculum. Here we examined to what extent participants were aware of their musical problem. To the question “Do you think that you lack a sense of music? », 20.3% of the controls said yes while 66.7 % of the pitch-based amusics did so, χ^2^(1)= 309.52, p < .001. Although we found the typical high rate of false alarms among controls,^20^ we also found in comparison a good proportion of self-disclosure among amusics (two-thirds of them).

### Family history of amusia

Given that the large majority of amusic participants appear aware of their deficit, they may be able to assess the family incidence of the disorder with fair accuracy. According to their report, 46% of their first-degree relatives were similarly affected (see Figure 3). This proportion was close to the 39% obtained by objective testing in the family aggregation study.^8^

**Figure 3.**
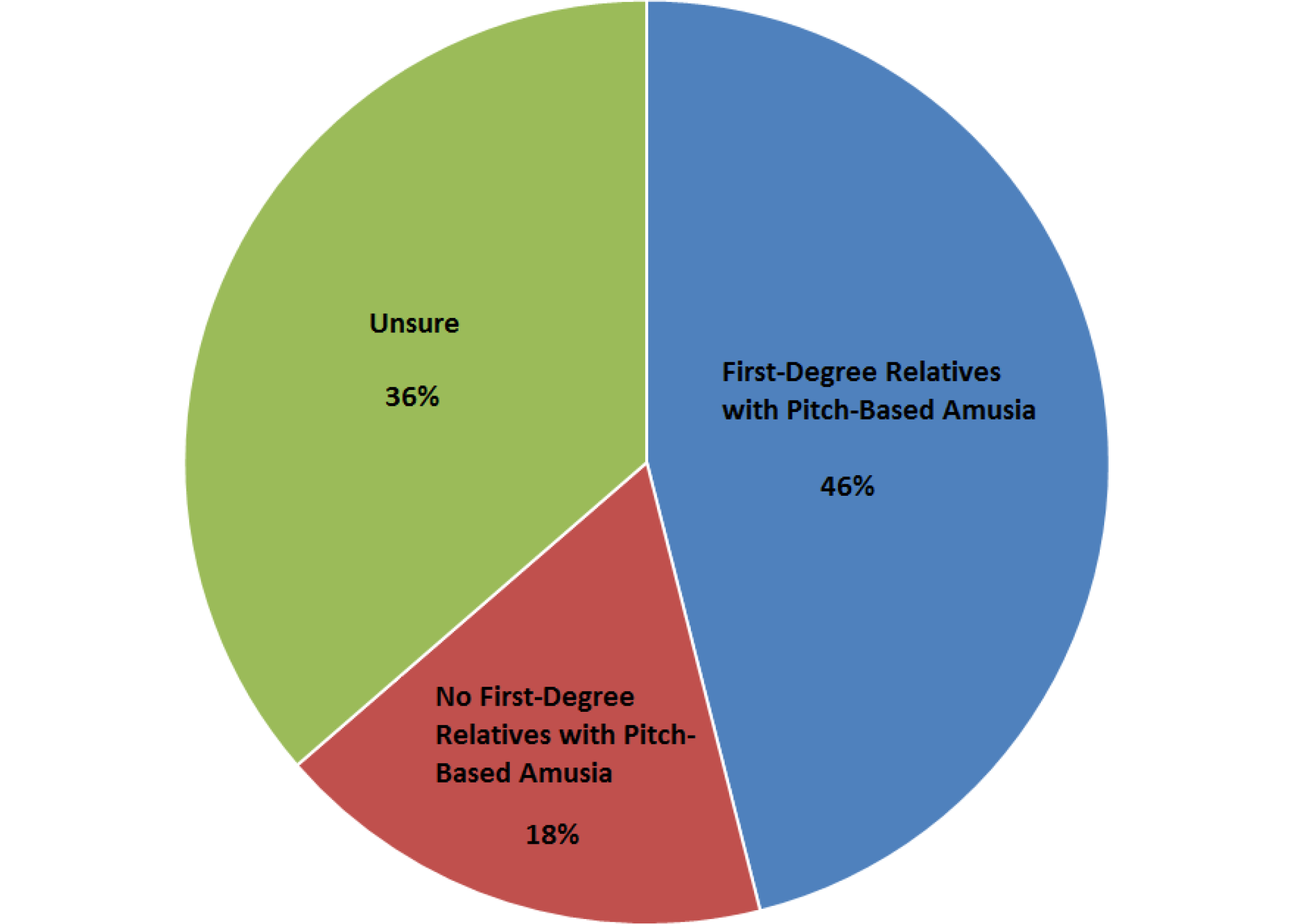
Heritability of congenital amusia by report.

### Co-morbidity: Associated disorders

There is no evidence that pitch-based amusia is associated to another neurodevelopmental disorder, besides problems with spatial orientation, χ^2^(1) = 6.28, *p* = .01 (Table 3). In contrast, a deficit in the off-beat test seems associated to many other neurodevelopmental disorders.

**Table 3.**
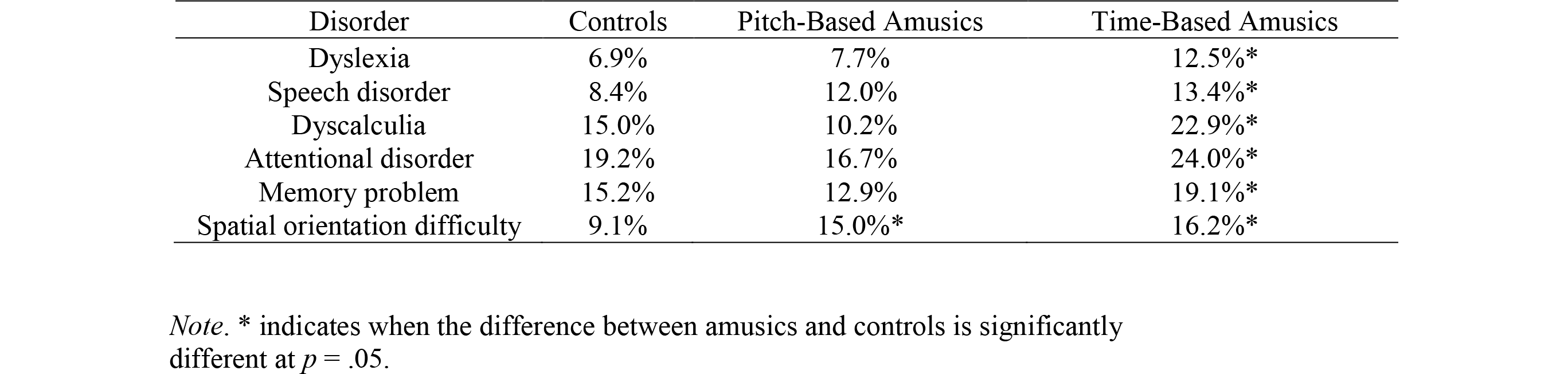
Association between pitch-based amusia, time-based amusia, and various neurodevelopmental disorders

## DISCUSSION

Here we present the first large scale (n = 16,625) study of congenital amusia that does not rely on self-referral and uses instead objective auditory tests. With three web-based tests, we obtained data over 10 years of testing from a large population with diverse demographic profiles. On this basis, we establish that congenital amusia affects 1.5% of the population with no marked difference between male and female participants.

To reach this prevalence of 1.5%, we used a conservative criterion for congenital amusia. Specifically, we considered participants to be amusic if they had an abnormal score (two standard deviations below the mean) on two separate tests requiring the detection of an out-of-key note, and a normal score on a test requiring the detection of an out-of-time note in the same unfamiliar melodies. Both the Scale and Off-key tests require access to tonal knowledge while the Scale test additionally entails a memory-based comparison process.

Therefore, an abnormal score in each test ascertains the genuineness of a deficit in detecting melodic key violation. On the other hand, the Off-key and control Off-beat tests have the same task demands but differ in the musical dimension that is measured (pitch vs. time). Therefore, a normal score on the control task excludes non-pitch-related difficulties with the testing situation. If we consider instead the more lenient but most widespread criterion for identification of congenital amusia, which corresponds to an abnormal score on the Scale test only, the prevalence of congenital amusia raises to 4.2%. The latter prevalence is in line with a prior large survey (n=1,000) aiming at a more homogenous Canadian population with a university level.^21^

Prevalence is determined statistically and depends on the selectivity of the tests to the manifestation of the disorder (phenotype). Prevalence is always relative. Its usefulness resides in its capacity to reflect the performance of a large population without confounding factors, such as attention and motivation. Interestingly, the disorder appears specific to the musical domain. The participants who are identified as amusic do not report problems with any other domain, such as language, besides spatial difficulties. The latter association between amusia and self-declared spatial difficulties has been observed previously.^19^ Moreover, there is some support for a possible link between amusia and visuo-spatial tasks, either in its congenital form^22,23^ or as a consequence of stroke.^24^ In the normal brain, pitch can be mapped to a vertical representation of space.^25^ Although this association between music and visuo-spatial skills is unlikely to underlie the majority of amusics’ cognitive difficulties with music perception it is an interesting avenue for exploring its neurobiological origins. Arguably, progress in the neurobiology of amusia requires a clarification of its relationship to other disorders.

In this regard, the other form of congenital amusia related to time (beat deafness) and not pitch might be more informative. Indeed, a majority of those impaired at detecting when a melodic tone was off-time (but normal at detecting when it was off-key) reported other developmental disorders such as dyscalculia and dyslexia. This finding calls for systematic cross-domain comparisons. In particular, the co-morbidity of beat deafness with dyslexia is predicted by recent research showing that children and adults who struggle to synchronize to a beat also struggle to read and have deficient neural encoding of sound.^26^ Reduced neural resources for temporal precision in both music and speech may result from a (common) genetic mutation. In contrast, the pitch-based form of congenital amusia would result from a distinct genetic etiology because it appears to emerge in isolation. These suggestions await objective testing. For instance, in the present study we did not test for the presence of facial recognition disorders (e,g., congenital prosopagnosia), which appear to have many neurobiological similarities with congenital amusia.^27^

Like the other neurodevelopmental disorders that are specific to a cognitive domain, such as selective language impairment (SLI), dyslexia, and congenital prosopagnosia, congenital amusia is heritable.^8^ Here, nearly half (46%) of the first-degree relatives are reported to be similarly affected. This proportion by report is close to the 39% obtained by test in our previous study^8^ and thus provides further empirical evidence across two population samples that there is a high familial risk related to congenital amusia. This presents a number of advantages not only for genetic association studies but also for the search of amusic participants. Using a general population sample is costly, particularly during childhood. Testing children at risk of developing amusia is much more efficient; for instance, in a general population sample with a rate of amusia at 1.5%, one would need to screen 1,000 children in order to obtain a sample of 15 amusic children; in a high-risk sample with a rate of amusia between 40-50%, testing less than 40 children will be required to yield the same number of affected cases.

In the search for the genetic variants of congenital amusia, it is useful to note that among the three tests used here the Scale test is to be preferred over the off-key and off-beat tests. According to a recent twin study,^28^ the Scale test yields predominantly additive genetic effects. The fact that musical training background was not assessed in this twin study is apparently not a serious concern. Here, we found little impact of musical training on the test scores. This may be due to the open nature of the instructions; depending on the context, an off-beat and off-key note can be acceptable for musically educated participants. Nevertheless, the small impact of musical education on the results should be treated cautiously. This factor may be confounded with socio-economic status, since in most countries extra-curricular musical lessons are not free and require dedication from the parents. In other words, a rich musical environment may not moderate the expression of congenital amusia but an impoverished musical environment may exacerbate a similar musical problem.

The next step in the genetic analysis of congenital amusia is to identify the specific genes involved and to relate these genes to the neuroanatomical anomalies found in the amusic brain. Importantly, genes do not specify cognitive functions but influence brain development. The amusic brain is characterized by impoverished communication in the network involving the inferior frontal cortex (BA 47) and the auditory cortex (BA 22) on the right side.^5–7^ Compared to controls, amusics have less white matter in the right inferior frontal cortex,^6^ while they have thicker cortex in the same right inferior frontal area and the right auditory area.^4^ These anomalies point to abnormal connectivity, supported by reduced fiber tracts in the arcuate fasciculus, due to anomalous neuronal migration or proliferation. Such malformations during cortical development are coded by these yet-to-be identified genes.^1^

In conclusion, congenital amusia is likely to be influenced by several genes that interact, both with each other and with the environment, to produce an overall susceptibility to the development of the disorder (i.e., a complex disorder). Its clear-cut behavioral expression (phenotype), is easily assessed by three auditory tests and its high heritability make the search for the responsible genes within reach.

## Supplementary Information

Supplementary information is available at (to be determined)

## References

1 Gingras B, Honing H, Peretz I,Trainor LJ, Fisher SE. Defining the biological bases of individual differences in musicality. Philos Trans R Soc Lond B Biol Sci 2015; 370:20140092.

2 Deriziotis P, Fisher SE. Neurogenomics of speech and language disorders: the road ahead. Genome Biol 2013; 14: 204.

3 Hyde KL, Zatorre RJ, Griffiths TD, Lerch JP, Peretz I. Morphometry of the amusic brain: A two-site study. Brain 2006; 129: 2562–2570.

4 Hyde KL, Lerch JP, Zatorre RJ, Griffiths TD, Evans AC, Peretz I. Cortical thickness in congenital amusia: when less is better than more. J Neurosci 2007; 27: 13028–13032.

5 Hyde KL, Zatorre RJ, Peretz I. Functional MRI evidence of an abnormal neural network for pitch processing in congenital amusia. Cereb Cortex 2011; 21: 292–299.

6 Loui P, Alsop D, Schlaug G. Tone deafness: a new disconnection syndrome? J Neurosci 2009; 29: 10215–10220.

7 Albouy P, Mattout J, Bouet R, Maby E, Sanchez G, Aguera PE et al. Impaired pitch perception and memory in congenital amusia: The deficit starts in the auditory cortex. Brain 2013; 136: 1639–1661.

8 Peretz I, Cummings S, Dubé M-P. The Genetics of Congenital Amusia (Tone Deafness): A Family-Aggregation Study. 2007 doi:10.1086/521337.

9 Peretz I, Kolinski R, Tramo M, Labrecque R, Hublet C, Demeurisse G et al. Functional Dissociations Following Bilateral Lesions Of Auditory-Cortex. Brain 1994; 117: 1283–1301.

10 Ayotte J, Peretz I, Hyde K. Congenital amusia: a group study of adults afflicted with a music-specific disorder. Brain 2002; 125: 238–251.

11 Zendel BR, Lagrois M-E, Robitaille N, Peretz I. Attending to Pitch Information Inhibits Processing of Pitch Information: The Curious Case of Amusia. J Neurosci 2015; 35: 3815–3824.

12 Drayna D, Manichaikul A, de Lange M, Snieder H, Spector T. Genetic correlates of musical pitch recognition in humans. Science 2001; 291: 1969–1972.

13 Stromswold K. Genetics of spoken language disorders. Hum Biol 1998; 70: 297–324.

14 Kalmus H, Fry DB. On tune deafness (dysmelodia): frequency, development, genetics and musical background. Ann Hum Genet 1980; 43: 369–382.

15 Peretz I, Champod AS, Hyde K. Varieties of Musical Disorders: The Montreal Battery of Evaluation of Amusia. In: Annals of the New York Academy of Sciences. 2003, pp 58–75.

16 Liu F, Patel AD, Fourcin A, Stewart L. Intonation processing in congenital amusia: Discrimination, identification and imitation. Brain 2010; 133: 1682–1693.

17 Phillips-Silver J, Toiviainen P, Gosselin N, Piché O, Nozaradan S, Palmer C et al. Born to dance but beat deaf: A new form of congenital amusia. Neuropsychologia 2011; 49: 961–969.

18 Wong PCM, Ciocca V, Chan AHD, Ha LYY, Tan LH, Peretz I. Effects of culture on musical pitch perception. PLoS One 2012; 7. doi:10.1371/journal.pone.0033424.

19 Peretz I, Peretz I, Gosselin N, Gosselin N, Tillman B, Tillman B et al. On-line identification of congenital amusia. Music Percept An Interdiscip J 2008; 25: 331–343

20 Cuddy LL, Balkwill LL, Peretz I, Holden RR. Musical difficulties are rare: a study of ‘tone deafness’ among university students. Ann N Y Acad Sci 2005; 1060311–324.

21 Provost M. The prevalence of congenital amusia. Master’s Thesis, Univ Montréal 2010.

22 Williamson VJ, Cocchini G, Stewart L. The relationship between pitch and space in congenital amusia. Brain Cogn 2011; 76: 70–76.

23 Tillmann B, Jolicoeur P, Ishihara M, Gosselin N, Bertrand O, Rossetti Y et al. The amusic brain: Lost in music, but not in space. PLoS One 2010; 5. doi:10.1371/journal.pone.0010173.

24 Särkämö T, Tervaniemi M, Soinila S, Autti T, Silvennoinen HM, Laine M et al. Cognitive deficits associated with acquired amusia after stroke: A neuropsychological follow-up study. Neuropsychologia 2009; 47: 2642–2651.

25 Lidji P, Kolinsky R, Lochy A, Morais J. Spatial associations for musical stimuli: a piano in the head? J Exp Psychol Hum Percept Perform 2007; 33: 1189–1207.

26 Carr KW, White-Schwoch T, Tierney AT, Strait DL, Kraus N. Beat synchronization predicts neural speech encoding and reading readiness in preschoolers. Proc Natl Acad Sci U S A 2014; 111: 14559–14564.

27 Avidan G, Behrmann M. Functional MRI Reveals Compromised Neural Integrity of the Face Processing Network in Congenital Prosopagnosia. Curr Biol 2009; 19: 1146–1150.

28 Seesjärvi E, Särkämö T, Vuoksimaa E, Tervaniemi M, Peretz I, Kaprio J. The Nature and Nurture of Melody: A Twin Study of Musical Pitch and Rhythm Perception. Behav Genet 2016; 46: 506–515.

